# Using EEG to understand how our brain elaborate information in stated choice experiments: Easy versus hard tasks in the choice of vehicles

**DOI:** 10.1101/2020.01.29.926162

**Authors:** Elisabetta Cherchi, Quoc C Vuong, Antonia Stergiou

## Abstract

In the current study, we aim to provide preliminary evidence that complex consumer choices depends on cognitive processes and executive functions that may not be fully captured by current stated choice (SC) approaches. To address this gap, here we combine the standard SC experiment with electroencephalogram (EEG) recordings while manipulating the cognitive demands of the task. Our study is applied to the choice context of a car purchase between a petrol and an electric vehicle. Respondents were asked to fill in a stated choice experiment online and a subsample of these respondents were then invited to participate in an EEG study during which they repeated the same SC experiment while we continuously recorded EEG signals from their scalp. We then modelled people’s choice behaviours in easy and hard decisions, and compared this analysis of their choice behaviour to their EEG responses in these two conditions. Our results confirm that hard decisions lead to higher cognitive demands and larger EEG responses in electrodes on the frontal part of the scalp and these demands can lead to choices inconsistent with the compensatory assumptions.

## INTRODUCTION

A persistent problem in using Stated Choice (SC) methods is that respondents often adopt decision processes that deviate from the assumption that individuals evaluate all the attributes presented in a compensatory way. As recently summarized by Cherchi and Hensher (2015), when respondents are “presented with a complex task, it is likely that they show disengagement, adopting simplifying strategies to reduce the mental effort required solving the problem. On the other hand, simplified survey tasks can be seemingly perceived as unrealistic by the respondents, leading to problems with respondents’ engagement, or respondents choosing based on other attributes not included in the design.” A body of literature has tried to incorporate different decision heuristics in the demand models, but this has often led to indistinguishable effects (see González-Valdés and Ortúzar, 2018). The problem is partly due to the theoretical assumptions (compensatory behaviour), and partly to the data and the information available. A growing literature on SC experiments has then focused on the problem of the decision process. Evidence, mainly based on model estimation results, shows that the attribute processing strategies adopted by respondents depends not only on the number of attributes and their levels but also on the importance and relevance of the attributes presented. If one attribute is much less important than the others to a respondent or its levels do not vary over a range that matters enough to result in a trade-off for the respondent, the attribute can be excluded from the decision process.

In an attempt to better understand how individuals make decisions, many researchers have recently used eye tracking technology to identify which visual information respondents pay attention to when making decisions (e.g., Balcombe et al., 2014, Krucien et al., 2014; Cherchi and Raja (2016); Uggeldahl et al., 2016; Meißner et al., 2016; Cherchi, 2018; Yang et al., 2015). This is a promising area of research, but at the moment, these papers mostly look at different effects of visual attention on the decision process. Eye movements can be a window into the human decision process by measuring what visual information people fixate on (e.g., the price of a car), and making a critical assumption that they are processing that information to make their decision (Koop and Johnson, 2011; Gidlöf et al, 2014; Itthipuripat, et al., 2015).

However, the critical assumption of eye movements do not always hold as people can attend and process visual information that they are not currently fixating on. Moreover making complex decisions involves executive functions such as working memory, inhibition and cognitive flexibility (Diamond, 2013) that are not captured by eye-movement data by themselves. For example, when given various attributes about two different types of cars (e.g., cost, CO_2_ emissions), people need to be able to hold and manipulate those attributes in working memory, retrieve relevant long-term information and possibly inhibit irrelevant information. A better understanding of these functions may be revealed using electroencephalography (EEG). This technique measures electrical potentials from the scalp with very high temporal resolution. The potentials reflect activation of different brain structures, which are known to be involved in executive functions, such as regions of the frontal or parietal cortex (Voigt et al., 2019). EEG measures in transport studies have been used mainly to study driving behaviour using driving simulators (e.g., Roman et al., 2001; Hernández et al., 2018; Park et al., 2018), and in some cases to study location functions and happiness in real environment to study (Mavros et al., 2016; Tilley et al., 2017). EEG has also been used to predict consumer choices and preferences (Avinash et al., 2018; Hakim & Levy, 2019; Golnar-Nik et al., 2019; Khushaba et al., 2012). In these studies, the researchers focus on EEG power in different frequency oscillation bands. These bands can be used to measure the involvement of some of the executive functions that play an important role in decision making (Ward, 2003); for example, power in the 4 to 8 Hz oscillation range (theta) reflects memory processes whereas power in the 8 to 12 Hz range (alpha) reflects attentional processes. Higher oscillation frequencies reflects the binding of sensory and cognitive information (Spitzer and Haegens, 2017).

In the current study, we aim to provide preliminary evidence that complex consumer choices depends on cognitive processes and executive functions (Diamond, 2013) that may not be fully captured by current SC approaches. To address this gap, here we combine the standard SC experiment with EEG recordings while manipulating the cognitive demands of the task. Our study is applied to the choice context of a car purchase between a petrol and an electric vehicle. Respondents from a panel provider were asked to fill in a stated choice experiment online and a subsample of these respondents were then invited to participate in an EEG study during which they repeated the same SC experiment while we continuously recorded EEG signals from their scalp. One strong assumption in the SC research, based on economics principles, is that respondents evaluate all the attributes presented, weigh them, and compute an overall benefit for each alternative they are given. Their stated choice is then predicted to be based on these benefits. According to these models, choice behaviours would not depend on the cognitive demands of the task. However, from a cognitive point of view, the stages in the decision process can be very demanding for complex choices such as choosing a type of car because there are many different attributes to weigh when trying to decide which type to purchase. We therefore first manipulated the difficulty of the choice scenarios to manipulate cognitive demands. We then modelled people’s choice behaviours in easy and hard decisions, and compared this analysis of their choice behaviour to their EEG responses in these two conditions. Based on previous work, we predict that hard decisions would lead to higher cognitive demands and larger EEG responses in electrodes on the frontal part of the scalp (Avinash et al., 2018; Hakim & Levy, 2019; Golnar-Nik et al., 2019; Khushaba et al., 2012). These demands can lead to choices inconsistent with the compensatory assumptions.

## DATA COLLECTION METHODOLOGY

### Stated choice experiment

As mentioned before, the goal of this study was to investigate brain activity, as measured by EEG power, while respondent were engaged in a stated choice experiment and making decisions. For this reason, we chose to use a relatively standard SC experiment, built upon previous works done to study individuals’ preferences towards an electric vehicle (see Jensen et al., 2014; Cherchi, 2017 and 2018). Our SC experiment consisted of a binary choice between an electric vehicle (EV) and an internal combustion vehicle (ICV) with the addition of a “neither of them” option. Five attributes were chosen to characterise the alternatives presented. These were purchase price, driving costs, driving performance (range), environmental effects (CO_2_ emissions) and the EV market share. The first four attributes represent the most significant attributes in the choice of EVs, according to the vast literature on EV. The last attribute, the EV market share, is not common in EV studies nor in SC experiments. It is taken from Cherchi (2017) and is a measure of descriptive norms, i.e., the influence that the action (or choice) of other people has on the individual’s choice. Information about the recharging station in UK was provided before starting the SC experiment, along with a link to the UK official webpage on the available network. Respondents were also carefully selected to guarantee that the information provided were realistic for them.

The levels of all the attributes were pivoted around the actual values of purchasing prices, driving costs, ranges and emissions of the UK car market, as well as the current market penetration of EVs. Following some pilot testing, for the EV market share we decided to include both the total number of new EV registration in 2019 in the UK and the percentage of new EV registered compared to the total number of cars registered in the same year. This differed from our previous work (Cherchi, 2017). We also decided to manipulate this information to give more emphasis to the low and high penetration level. The attribute had 3 levels. The lower level was set to 4,988 new EV registrations that corresponded to the current EV market in the UK. Other than the absolute level, we also provided information about the corresponding market share manipulated as follows: “still only 3% of the Market Share”. The mid-level was set equal to 16,627 new EV registrations but the market share information (10% of the market share) was not manipulated. The higher level was set to 49,880 EV new registrations and the market share information was manipulated as follows: “a record 30% of the Market Share”.

The SC experiment was customised based on the car size that respondents intended to buy within the next 5 or 10 years or the last car purchased in the household. In order to ensure that the prices range displayed in the SC experiment was realistic, respondents were also asked to indicate the range of prices for their next or past purchase. Three ranges were defined, corresponding to a small, medium or large car. Some screening information was collected to guarantee realism. In particular, respondents needed to have a drive licence, and they had to live in an area where it is realistic to install a recharging station at home.

Figure 1 shows an example of a choice scenario that was used for both the online survey and the EEG study. Differently from what was reported in Cherchi (2017), we did not need to simplify the information reported because the amount of information do not affect the EEG analysis.

**Figure 1:**
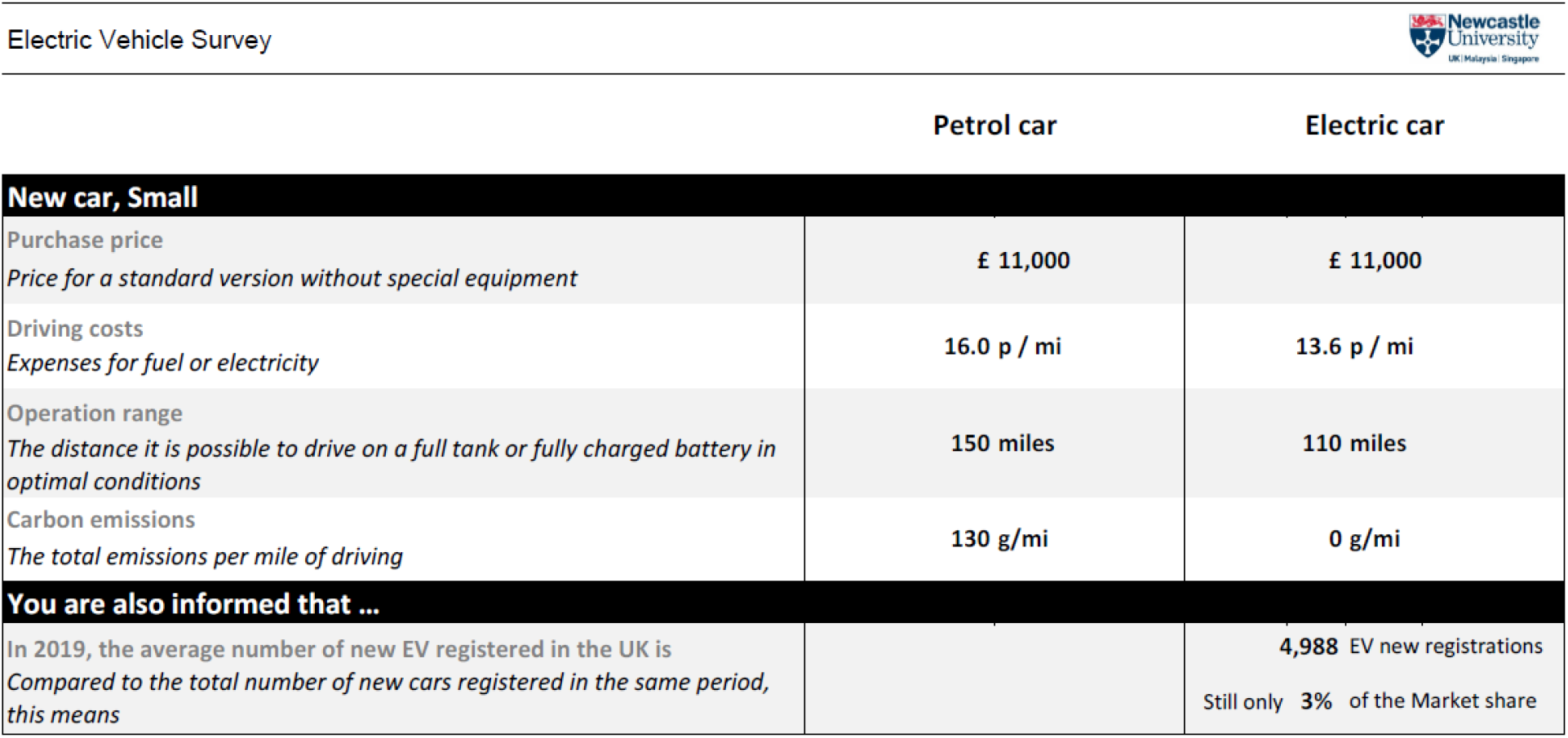
Example of an SC scenario used in the current study

An orthogonal design was built, accounting only for main effects, for a total of 16 choice scenarios. Using priors available from previous studies, the scenarios were built and simulated scenarios were built in a way to ensure that roughly in 8 scenarios the probabilities to choose EV and ICV were within 39% and 59% (this happens when the level of the attributes is similar between alternatives) and in the other 8 scenarios were higher that 59% or lower than 39%. When the alternatives have a similar probability to be chosen, the decision is expected to be difficult. We defined these cases as “hard” scenarios. When the difference in the probability of choosing the alternatives is large, it means that one alternative is much better than the other, and so the choice is expected to be simpler. We defined these cases as “easy” scenarios. It is important to mention that these easy scenarios do not include cases with dominant alternatives (i.e., an alternative with choice probability close to 100%). That is, for easy scenarios a trade-off between the two alternatives is still possible.

The 16 scenarios were randomly divided into 2 blocks of 8 scenarios presented to each respondent. In each block, we randomised the order of the scenarios presented and the position (left or right) of the two alternatives (EV or ICV).

### Online Survey

The online survey was implemented in SurveyEngine, with the kind assistance from the team at SurveyEngine GmbH in Berlin. Other than the SC experiment, the survey included several questions about the number and type (size and type of engine) of cars available in the household, the specific car most used by the respondent, the purpose this car was most used for, and the kilometres driven daily. We then asked whether (and which car exactly) they intended to replace or to buy as a new car. This information was used to customise the SC scenarios presented to that participant. After the SC experiment, we asked additional questions related to socio-economic characteristics and where they live (in particular if they had the possibility to install an electric charge at home). Finally, a set of attitudinal questions was asked, measuring attitudes toward environment and injunctive norms. The statements were taken from Cherchi (2017).

A sample of 118 participants was randomly selected from members of a panel, trying also to match the gender, age and education balance. Participants were randomly grouped into lots of 20 participants. A £20 voucher was awarded to a randomly selected individual for each lot.

### EEG Study

#### Description of the EEG recordings

Participants completed the same SC experiment in the laboratory while we recorded EEG responses. The EEG recordings were acquired using a 128-electrode HydroCel Geodesic Sensor Net (Electrical Geodesic Inc., Eugene, OR) and amplified using an EGI Net Amps 400 amplifier. The sampling rate was 1000 Hz and the data was referenced on-line to the vertex electrode (Cz). We checked that the impedance at each electrode was less than 50 KΩ before starting the continuous EEG recording.

#### Description of the protocol

Twenty participants from the larger sample consented to participate in the EEG follow-up experiment after completing the online survey (15 females; mean age: 39.6 years; range: 20 to 63 years). Ethics was approved by the Newcastle Faculty of Medical Sciences ethics review panel; all participants signed a consent form. Participants were reimbursed £30 for travel costs to the EEG lab and for their time.

There were 16 SC scenarios for the three different preferred car size (small, medium and large). These scenarios included the eight from the online survey and eight new ones. For the small car size, there were 8 easy and 8 hard scenarios; for the medium size, there were 9 easy and 7 hard scenarios; and for the large size, there were 11 easy and 5 hard scenarios.

Participants complete two blocks of the SC task, with a break between blocks. The 16 scenarios for each participant’s preferred car size were presented in a random order on each block for a total of 32 trials. This ensured that there were a sufficient number of trials in each condition for a high signal-to-noise ratio in the EEG data. Approximately half the participants received information about the EV on the left column and half on the right (matched to how the information was presented to that participant on the online survey). They sat approximately 50 cm from the computer monitor. On each trial, they were shown a white fixation cross on a grey background for 1 sec, followed by a 0.5 sec grey screen, followed by a choice scenario for 35 sec. The scenario was then replaced by a blank grey screen for 1 sec, followed by a grey response screen with the three possible responses indicated in white text (“electric”, “neither”, “petrol”). Participants responded by pressing the arrow key corresponding to their choice on the numeric keypad with the right hand. There was a 2-sec grey screen following the response before the next trial began. The experiment was synchronised to the EEG recording by sending an event marker on the onset of each choice scenario. The event marker coded the condition (easy or hard).

### DATA ANALYSIS AND RESULTS

#### EEG results

The EEG data were analysed off-line using EEGLAB (version 2019; Delorme & Makeig, 2004). For each participant, the data were resampled to 250 Hz (to speed up analyses), band-passed filtered with frequency cut-offs of 0.3 and 30 Hz, and segmented into 25-sec easy and hard epochs relative to the onset of the choice scenario (32 epochs total). Next, an independent component analysis (ICA) was run on the epoched data separately for each condition, and components related to eye blinks, eye movements and muscle artefacts were manually rejected.

A fast Fourier transform (FFT) was used to compute the power spectral density (PSD) for each electrode and epoch on the pre-processed and cleaned EEG data. The PSD reflects the power at each frequency between 0.3 and 30 Hz. The median PSD was calculated across epochs separately for each condition to allow for comparisons between easy and hard choice scenarios. For each condition, the power was averaged within four frequency bands: delta (1 to 3 Hz), theta (4 to 7 Hz), alpha (8 to 12 Hz) and beta (13 to 30 Hz). Following previous studies (Avinash et al., 2018; Hakim & Levy, 2019; Golnar-Nik et al., 2019; Khushaba et al., 2012), we focused on frontal, central and parietal clusters of electrodes in the left and right hemispheres. For each cluster, we averaged the power across the electrodes in that region. The power data were submitted to analyses of variance (ANOVAs) with the factors: cluster, frequency band and condition. For all statistical analyses, an alpha = .05 was used as the significance level and partial-eta-squared as a measure of effect size (0.06 to 0.14 considered medium effect size, and > .14 considered large effect size).

Figure 2 shows the mean EEG power (averaged across participants) for each frequency band, cluster and condition. Most of the power in all clusters is concentrated in the slow delta band, and power was highest in frontal clusters. We submitted the power data to a separate ANOVA for each frequency with cluster (frontal, central, parietal; averaged across hemisphere) and condition (easy, hard) as repeated measures. There was a large main effect of cluster for all frequency bands (delta: F(2,38) = 17.83, p < .001, partial-eta-squared = .48; theta: F(2,38) = 21.57, p < .001, partial-eta-squared = .53; alpha: F(2,38) = 24.93, p < .001, partial-eta-squared = .57; beta: F(2,38) = 43.50, p < .001, partial-eta-squared = .70). The condition factor was marginally significant for the slower delta (F(1,19) = 3.52, p = .08, partial-eta-square = .16) and theta (F(1,19) = 3.20, p = .09, partial-eta-square = .14) bands, and it was significant for the faster alpha (F(1,19) = 4.91, p = .04, partial-eta-squared = .21) and beta (F(1,19) = 7.99, p = .01, partial-eta-squared = .30) bands. Previous studies suggested that power in frontal theta band could be used to predict consumer decisions and preferences, as frontal brain regions are thought to be involved in complex decision making (Avinash et al., 2018; Hakim & Levy, 2019; Golnar-Nik et al., 2019; Khushaba et al., 2012). In line with these studies, there was some indication that power differences between trial types were more enhanced for frontal compared to the other electrode clusters for the theta (the interaction between cluster and condition approached significance: F(2,38) = 2.02, p = .15, partial-eta-squared = .10) and beta (F(2,38) = 2.29, p = .12, partial-eta-squared = .11) bands.

**Figure 2.**
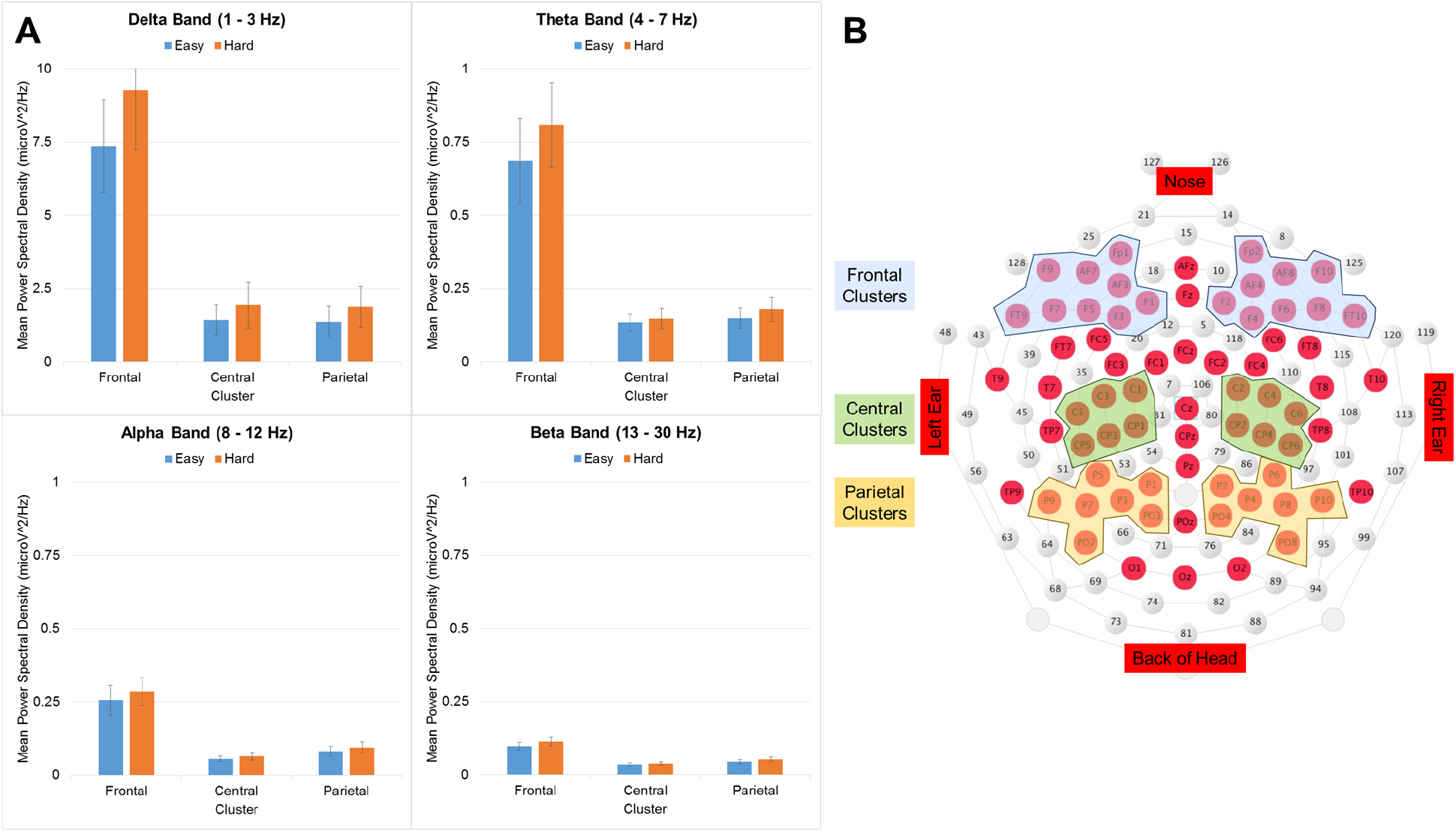
(A) Results from the EEG study. (B) Location of the electrode clusters.

Our preliminary results suggest that more difficult decisions can recruit higher oscillation frequencies (e.g., beta) which may play a role in helping people bind information (Spitzer and Haegens, 2017; Ward, 2003; although we did not analysed oscillation frequencies beyond 30 Hz). Previous work showed that theta power in frontal electrodes play an important role in reflecting consumers’ preferences for a product (e.g., Golnir-Nik et al., 2019). Although not significant in the current study, we found a similar trend that frontal theta power can reflect consumers’ preferences when they had more than one alternative to choose from, which require them to weigh the attributes across both alternatives. Further investigation is needed as some of the findings approached significance and effect sizes were in the medium to large range (partial-eta-squared > 0.06). Furthermore, electrical potentials measured at any one position on the scalp reflect summed activities across several cortical regions so the cortical sources of EEG signals cannot be resolved exactly. These sources can be estimated from EEG signals (Pascual-Marqui, 2002) but future studies can use a similar SC experiment with functional magnetic resonance imaging to more accurately localise the specific brain regions involved.

#### Analyses of the discrete choices

Figure 3 reports the distribution of the choices from the sample online and the sample who did the SC experiment in the lab with the EEG. Table 2 reports the % of scenarios in the SC experiments where the simulated probability was lower than 40% or higher than 60% (easy scenarios). For the hard scenarios, i.e. where the probability is within 40%-60%, the probability of choosing EV or ICV is 50-50%. We note that in our SC experiment, the EV was slightly favoured for small and medium car size, and vice versa for the large car size. However, this is not relevant given that one of the main objectives of this research was to analyse EEG power while participants were making choices.

**Table 2:** Percentage of easy scenarios in the SC experiment

| Car Class | Petrol | EV | None |
| --- | --- | --- | --- |
| small | 33% | 67% | 0 |
| medium | 40% | 60% | 0 |
| large | 67% | 33% | 0 |

**Figure 3:**
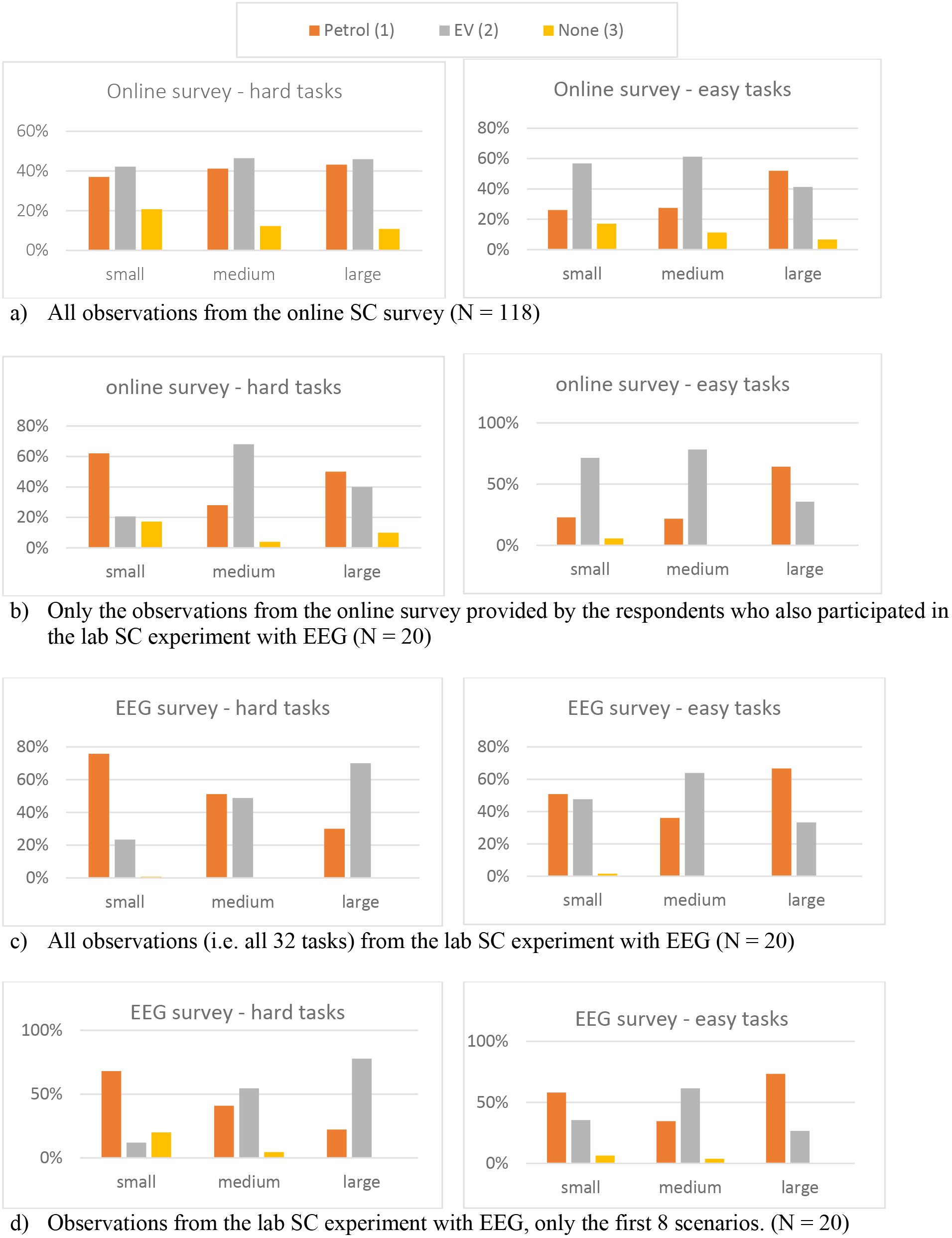
Distribution of the choices from the SC experiments

First we note that the distribution of the choices in the online sample reflect the distribution assumed in the SC experiment. The choices split almost equally between the EV and ICV, for all car classes, when the tasks were hard, while there is a marked preference for EV for small and medium cars (and for ICV for large car) when the tasks were easy.

Interestingly, this is not the case when respondents performed the same experiment in the lab with EEG. In particular, results are opposite for the easy tasks, small cars, where respondents prefer ICV more than EV cars. The case with hard tasks, however, is the one that shows the most striking difference, with respondents clearly preferring ICV for small cars and EV for large cars. We note that this effect is the same if we consider only the first 8 choice scenarios presented, or all 32 scenarios. We can safely rule out the assumption of fatigue, learning or practise effect in the analysis of these results (recall that in the online survey, each respondent evaluate 8 choice scenarios, while in the lab experiment 32). Finally, we note the number of time the alternative “none” was chosen in the lab experiment is significantly lower than in the online experiment.

#### Preferences estimation based on compensatory assumptions

Table 1 reports the results of the mixed logit models estimated using the stated choice data collected online and the stated choice data collected during the EEG experiment. The model structure used in this paper is a mixed logit model typically used to model choices among a set of discrete and mutually exclusive alternatives. Mixed logit models are grounded on the concept of rationality that assumes that individuals possess a mental order of preferences that allow them to have perfect information about all the available options and the possible consequences of their actions. They are able then to associate to each option a utility function that measures the level of satisfaction they derive from it and to make finally their choice coherently with their preferences and with the constrains upon them. The concept of utility, i.e. a unique index that summarises the level of satisfaction received from the eventual choice of each alternative, implicitly assumes the concept of trade-off among attributes, i.e. that a bad attribute can be compensated by a good attributes.

In the general case, where an individual *q* has to choose among *j* alternatives, in *t* different scenarios, the utility, in the mixed logit model, can be formalised as follows:

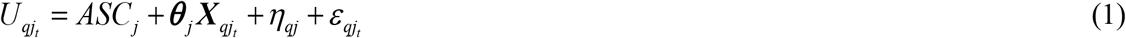

Where U_*qjt*_ is the utitliy that individual *q* associates to alternative *j* in the choice task *t*. ***X*** is a vector that includes all the attributes presented in the SC experiment and ***θ*** a vector of the associated coefficients. ASC is the alternative specific constant. η is an error term distributed Normal (0, σ^2^) that accounts for correlation among observations of the same individual and ε an error term identically and independently distributed extreme value type 1.

The conditional probability to choose the sequence of choices *j_t_*, is then given by the product over the SC choice tasks of multinomial logit probabilities conditional on the realisation of η. The unconditional choice probabilities is given by:

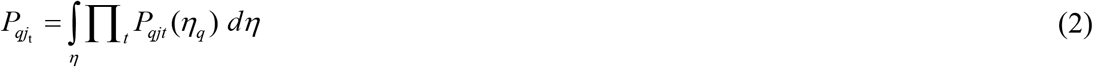

Models are estimated by maximum likelihood estimation, using PythonBiogeme (Bierlaire 2016).

Table 3 reports the results of the mixed logit models estimated using the stated choice data collected online. The first column (labelled “full experiment”) reports the results from the entire dataset, while the other two columns report the results specification estimated using only the tasks classified as “easy” and as “hard”. We can see that all coefficients have the expected sign, in agreement with the microeconomic theory. As expected, the purchase price is the most relevant attribute in the choice of the vehicle, followed by the range. In line with the literature on discrete choice models, panel effect (i.e. correlation among observations from the same individual, captured by the attributes called “sigma” and “rho”) is highly significant and reveals also the presence of significant random heterogeneity in the preference for EV and ICV.

**Table 3.** Model estimation results from the online survey

|  | Full experiment |  | Easy tasks |  | Hard tasks |  |
| --- | --- | --- | --- | --- | --- | --- |
|  | Value | Robust t-test. | Value | Robust t-test. | Value | Robust t-test. |
| ASC (EV) | 0.286 | 0.29 | 2.120 | 1.37 | 1.390 | 0.90 |
| ASC (ICV) | -9.36 | -2.48 | -3.110 | -1.08 | -2.690 | -1.13 |
| SIGMA (EV) | 5.78 | 4.04 | 5.380 | 2.89 | 3.330 | 2.54 |
| SIGMA (ICV) | 2.53 | 6.92 | 2.840 | 5.56 | 2.710 | 6.01 |
| RHO2 (ICV) | 6.22 | 4.59 | 5.640 | 3.22 | 4.050 | 3.95 |
| **Characteristics of the vehicles** |  |  |  |  |  |  |
| Purchase price [10,000 GBP] | -4.38 | -4.76 | -3.650 | -3.83 | -1.460 | -1.71 |
| Fuel costs [*pence/mile*] | -0.134 | -0.34 | -0.388 | -0.43 | 0.544 | 0.90 |
| CO2 Emissions [*grams/mile*] | -1.01 | -1.58 | -1.040 | -1.05 | 0.376 | 0.35 |
| Driving range (EV) [100 miles] | 3.64 | 6.47 | 6.000 | 5.90 | 1.260 | 1.13 |
| Driving range (ICV) [100 miles] | 2.22 | 3.72 | 4.490 | 4.17 | 0.499 | 0.55 |
| **Measures of Descriptive Norms** |  |  |  |  |  |  |
| EV new registration [10,000] | 0.0363 | 0.65 | -0.063 | -0.78 | -0.391 | -1.97 |
| **Overall statistics** |  |  |  |  |  |  |
| Number of draws | 1000 |  | 1000 |  | 1000 |  |
| Number of observations | 944 |  | 504 |  | 440 |  |
| Log-likelihood at the maximum | -623.757 |  | -338.36 |  | -352.50 |  |
| Akaike Information Criterion: | 1269.513 |  | 698.71 |  | 7226.99 |  |
| Bayesian Information Criterion: | 1322.865 |  | 745.16 |  | 771.95 |  |

The model estimation highlighted differences in participants’ choice behaviours between the easy and hard tasks. In the model estimated with only the easy tasks, purchase price and range became slightly more significant, and in particular the preference for the range double, while CO_2_ emission became slightly less significant and the descriptive norms takes a wrong sign (though not significant). Thus, it seemed that respondents were still engaged for easy decisions, but they had a tendency to focus more on few key attributes. The model estimated with only the hard tasks, on the other hand, revealed a clear deviation from the predicted compensatory behaviour. None of the attribute, not even the purchase price and the range, is significant at 95%. More analyses are required to identify if there are simplifying strategies behind these choices. At the moment, it seems that the decision process is almost random.

Overall the model estimation results suggest that participants were not equally using the same attribute values in the easy and hard tasks, and so additional factors may be involved. In line with this interpretation, our preliminary EEG results suggest that people may use executive functions differently for easy and hard decisions. That is, hard tasks, which we constructed to require more cognitive demands, may engage more executive functions as reflected by increased frontal power and increased binding and manipulation of information as reflected by increased power in the beta frequency band.

## CONCLUSIONS

Using a stated choice experiment built by manipulating the difficulty of the choice task, this work showed easy and hard tasks lead to significant differences in participants’ choice behaviours and in the EEG responses. Our results confirm that hard decisions lead to higher cognitive demands and larger EEG responses in electrodes on the frontal part of the scalp and these demands can lead to choices inconsistent with the compensatory assumptions. In the SC literature, it is recommended that the tasks should not be too easy, otherwise the choice would not be informative in terms of the trade-off between attributes; but not too complex, otherwise respondents may find the task too difficult and so their choices may not be based on trading-off the attributes. Both our behavioural and neural findings support this recommendation.

It is important to note that the definition of hard and easy tasks carry a certain degrees of arbitrariness. It depends on the actual levels presented and how similar respondents perceived those levels. Finally, it is important to stress that an easy choice does not necessarily imply that one alternative is dominant over the other. We did not have any dominant alternatives in our experimental design.

EEG can be a very effective tool to address issues in complex decision making but it has limitations. For example, we allowed participants to freely move their eyes to read the text, which can cause eye-movement related artefacts (ICA help reduce the impact of eye movement artifacts). The EEG signals are typically noisy so many trials are needed to increase the signal-to-noise ratio. This made it challenging to use EEG to study SC tasks, because of several behavioural implications related with the length of the experiment. Lastly, the brain sources generating the EEG signals measured on the scalp cannot be precisely localised so caution is needed when relating EEG power to brain activity. Despite these limitations, this is the first study to combine SC experiments with EEG results. The preliminary results are very promising, but more data are needed to reach more robust conclusions.

## AUTHOR CONTRIBUTIONS

The authors confirm that Elisabetta Cherchi and Quoc Vuong have equally contributed to all parts of the research and in writing the paper. Antonia Stergiou has been responsible for the lab experiment with EEG.

